# Correlation of receptor density and mRNA expression patterns in the human cerebral cortex

**DOI:** 10.1101/2021.12.23.473947

**Authors:** Murgaš Matej, Michenthaler Paul, Reed Murray Bruce, Gryglewski Gregor, Lanzenberger Rupert

## Abstract

Changes in distribution of associated molecular targets have been reported across several neuropsychiatric disorders. However, the high-resolution topology of most proteins is unknown and simultaneous *in vivo* measurement in multi-receptor systems is complicated. To account for the missing proteomic information, mRNA transcripts are typically used as a surrogate. Nonetheless, post-transcriptional and post-translational processes might cause the discrepancy between the final distribution of proteins and gene expression patterns. Therefore, this study aims to investigate *ex vivo* links between mRNA expression and corresponding receptor density in the human cerebral cortex.

To this end, autoradiography data on the density of 15 different receptors in 38 brain regions were correlated with the expression patterns of 50 associated genes derived from microarray data (mA), RNA sequencing data (RNA-Seq) provided by the Allen Human Brain Atlas and predicted mRNA expression patterns (pred-mRNA). Spearman’s rank correlation was used to evaluate the possible links between proteomic data and mRNA expression patterns.

Correlations between mRNA and protein density varied greatly between targets: Positive associations were found for e.g. the serotonin 1A (pred-mRNA: r_s_ = 0.708; mA: r_s_ = 0.601) or kainate receptor (pred-mRNA: r_s_ = 0.655; mA: r_s_ = 0.601; RNA-Seq: r_s_ = 0.575), while most of the investigated target receptors showed low or negative correlations.

The high variability in the correspondence of mRNA expression and receptor warrants caution when inferring the topology of molecular targets in the brain from transcriptome data. This highlights the longstanding value of molecular imaging data and the need for comprehensive proteomic data.

## 1. Introduction

To unravel underlying molecular mechanisms in the human brain, it is of paramount importance to establish an *in vivo* analysis method of multiple receptor systems in both healthy persons and patients. Currently*, in vivo* imaging modalities are usually limited to specific receptors; e.g positron emission tomography (PET), which necessitates the use of a radiotracer to allow for the investigation of very few targets. Moreover, systems utilizing two tracers are currently under development, whilst the number of targets concurrently detectable and quantifiable in PET will remain critically limited for the foreseeable future, as currently dual tracer PET is developed for a limited application (e.g. hepatocellular carcinoma) and remains prohibitively expensive (Filippi et al., 2019; Lu et al., 2019). Current studies suggest that it is highly unlikely that changes in a singular receptor system confer all symptoms of brain disorders as complex as, e.g. depression (Villas Boas et al., 2019). The numerous pharmacological targets of drugs used in the treatment of neuropsychiatric disorders further strengthen this claim. Furthermore, many drugs used in contemporary psychopharmacological treatment show efficacy in different psychiatric disorders, pointing towards shared mechanisms between them (Lamers et al., 2011; Rainer et al., 2012).

In recent years, neuroimaging techniques quantifying protein distribution (e.g. PET) have become commonplace in psychiatric research (Lui et al., 2016), e.g. to investigate possible effects of electroconvulsive therapy in depressed patients (Baldinger-Melich et al., 2019) or the effect of bright light therapy in patients with affective disorders (Spies et al., 2018). However some of the basic hypotheses of modern psychiatry, including the monoamine hypothesis, still stem from decade-old *post mortem* findings (Cosci and Chouinard, 2019; Horton, 1992). To this end, shedding light on associations between *ex vivo* data generated using radioligands to mark specific structures (such as receptors, enzymes, etc.) and their underlying mRNA expression patterns could pave the way toward more complete analysis and potentially allow for extrapolation of protein distributions based on molecular information.

Modern multimodal atlases like the Allen Human Brain Atlas (AHBA) provide valuable insights into the cerebral mRNA expression patterns and by this virtue might allow estimation of the respective protein receptor abundance and distribution. Previous studies have demonstrated similarities between mRNA expression patterns and the distribution of certain receptors, e.g. serotonin 1A and 2A (Beliveau et al., 2017; Komorowski et al., 2017; Unterholzner et al., 2020), dopamine D3 (Komorowski et al., 2020b), or enzymes (Lohith et al., 2017) measured using PET. Moreover, associations between mRNA expression patterns and task-specific activations measured by fMRI were revealed for reward and emotion processing(Komorowski et al., 2020a). Contrary to these findings, no relations were discovered between protein distribution and gene expression of monoamine oxidase A (Godbersen et al., 2021) and serotonin transporter (Komorowski et al., 2017). Preliminary investigations have also shown mixed results when comparing transcriptomic data to receptor densities measured by autoradiography (Murgas et al., 2020, 2021a, 2021b, 2021c).

Uncovering stable defining correlations across different brain regions, mRNA expression and resulting gene product abundance may enable future estimation of protein distribution based on initial gene expression patterns and potentially even support personalized treatments in psychiatry. Notwithstanding, a number of challenges remain as studies looking for associations between mRNA expression and protein distribution often fail to deliver significant positive results and even significant results are inconsistent in their findings (Arnatkevičiūtė et al., 2019). In the current study, we compared a collection of cortical multi-receptor proteomic autoradiography fingerprints measured by Jülich Research Centre (Zilles and Palomero-Gallagher, 2017) to a corresponding dataset of *ex vivo* gene expression patterns adopted from Allen Human Brain Atlas measures.

## 2. Materials and Methods

To elucidate possible links between molecular mechanisms and protein distributions, we compare the spatial distribution of 15 different receptors and mRNA transcripts associated with them or their subunits. In particular, we study three ionotropic glutamate receptors - AMPA, Kainate and NMDA; GABA receptors - GABA_A_, GABA_A/BZ_ (benzodiazepine binding site) and GABA_B_; cholinergic receptors - muscarinic M_1_, M_2_, M_3_ and nicotinic α_4_β_2_; adrenoceptors α_1_ and α_2_; serotonin receptors 5-HT_1A_ and 5-HT_2_; dopamine receptor D_1_. The full list of investigated relations is in Table 1.

**Table 1:** Overview of target receptors and their corresponding gene expression patterns associated with the receptor or its possible subunit.

| Receptor | Associated gene expressions |
| --- | --- |
| AMPA | *GRIA1, GRIA2, GRIA3, GRIA4* |
| Kainate | *GRIK1, GRIK2, GRIK3, GRIK4* |
| NMDA | *GRIN1, GRIN2A, GRIN2B, GRIN2C, GRIN2D, GRIN3A, GRIN3B* |
| GABA<sub>A</sub> | *GABRA1, GABRA2, GABRA3, GABRA4, GABRA5, GABRA6, GABRB1, GABRB2, GABRB3, GABRD, GABRE, GABRG1, GABRG2, GABRG3, GABRP, GABRQ* |
| GABA<sub>B</sub> | *GABBR1, GABBR2* |
| GABA<sub>A/BZ</sub> | *GABRA1, GABRA2, GABRA3, GABRA5, GABRB1, GABRB2, GABRB3, GABRD, GABRE, GABRG1, GABRG2, GABRG3, GABRP, GABRQ* |
| M<sub>1</sub> | *CHRM1* |
| M<sub>2</sub> | *CHRM2* |
| M<sub>3</sub> | *CHRM3* |
| $\alpha_4\beta_2$ | *CHRNA4, CHRNB2* |
| $\alpha_1$ | *ADRA1A, ADRA1B, ADRA1C* |
| $\alpha_2$ | *ADRA2A, ADRA2B, ADRA2C* |
| 5-HT<sub>1A</sub> | *HTR1A* |
| 5-HT<sub>2</sub> | *HTR2A, HTR2B, HTR2C* |
| D<sub>1</sub> | *DRD1* |

### 2.1 Receptor distribution

In this data-driven approach, we utilize a readily available dataset (Zilles and Palomero-Gallagher, 2017) comprising of autoradiography data (autoRTG) acquired from the brains of three healthy donors (72-77 years old, 2 males) as a source for multi-receptor profiles across 38 cortical areas defined in the original paper of the data. Regional values represent mean receptor density (fmol/mg) over all cortical layers.

### 2.2 Gene expression

Corresponding mRNA expression data are adopted from the AHBA (ALLEN Human Brain Atlas, 2013; Hawrylycz et al., 2012), which provide data on a variety of transcripts and a considerable number of sampling sites. The atlas contains microarray data (mA) from six adult donors (24-57 years old, 5 males) and additional RNA sequencing (RNA-Seq) data of two donors. To account for the spatially incomplete data of AHBA, we also make use of predicted mRNA expression patterns (pred-mRNA). Briefly, AHBA samples for microarray data are mirrored along the x-axis of MNI space to overcome a substantial difference in the number of samples from the left and right hemispheres in AHBA. Utilizing kriging via Gaussian process regression, continuous mRNA expression across the cortex is predicted (Gryglewski et al., 2018). Maps are publicly available at http://www.meduniwien.ac.at/neuroimaging/mRNA.html. pred-mRNA are further represented by mean regional values of 38 cytoarchitectonicaly distinct areas defined in Julich-Brain atlas (Amunts et al., 2020) (Figure 1, Suppl. Table 1).

**Figure 1:**
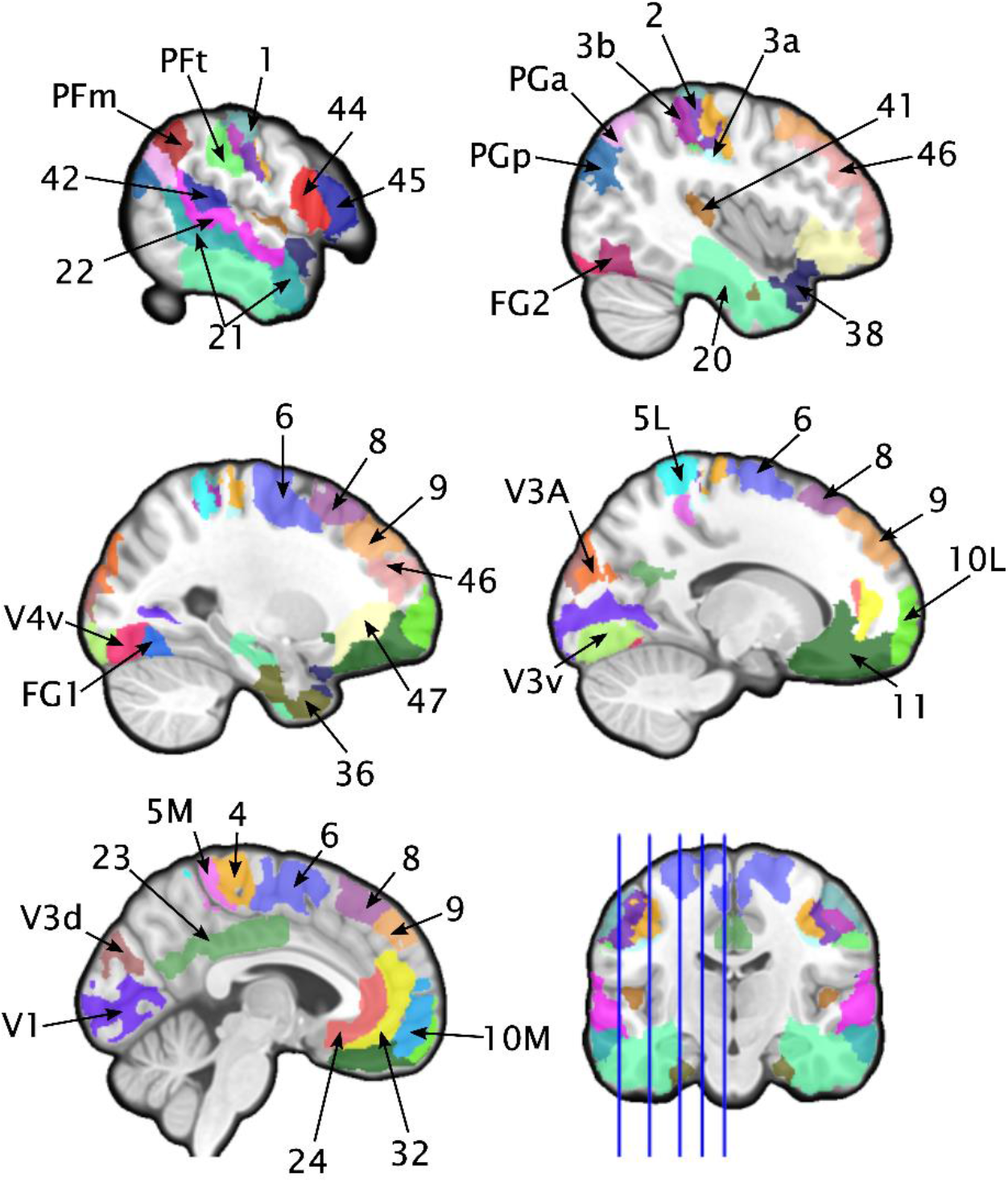
Visual representation of ROIs included in analysis. Diverse colors indicate covered areas. For the full list of the regions see supplementary table 1.

Microarray data from the AHBA data set consists of 3702 samples that need to be accordingly pre-processed (Arnatkevičiūtė et al., 2019). Therefore, we first identify probes that have a significantly different mean signal from the background and at the same time the difference between their background-subtracted signal and background is more than 2.6 (ALLEN Human Brain Atlas, 2013). As there are typically multiple probes associated with one target within one sample, the mean value is calculated for the particular sample. Next, samples are assigned to a particular region based on their spatial location. This is achieved by warping the Colin27 MNI template (Holmes et al., 1998) to the subjects T1 using SPM12 (The Wellcome Centre for Human Neuroimaging, www.fil.ion.ucl.ac.uk) to the individual space of each brain from AHBA separately. As a result, each of the 1390 microarray samples is assigned to 38 regions from the Julich-Brain atlas. To account for the inter-subject variability of AHBA mA data, scaled robust sigmoid normalization is applied (Fulcher et al., 2013). Subsequently, average regional values are calculated as the mean over the probe values, for each gene expression.

With the current trend in the RNA analysis directing to the more robust and reliable gene expression quantification (Wang et al., 2014), we also make use of two additional RNA-Seq data available from the AHBA (ALLEN Human Brain Atlas, 2013; Hawrylycz et al., 2012; Shen et al., 2012). The deformation fields from the previous step were additionally used to assign the 242 RNA sequencing samples to individual regions, resulting in 165 RNASeq samples associated with the selected regions. As not all of the selected areas are associated with the samples for RNA-Seq data only those ROIs with at least 1 independent data point reported were retained, resulting in 34 regions from RNASeq for analysis. Subsequently, all samples are averaged region-wise and represent the mean Transcripts Per Kilobase Million (TPM). To investigate the discrepancy between RNA-Seq data and the other 2 data set based on microarray data, we compared RNA-Seq data to pred-mRNA and mA.

### 2.3 Statistical analysis

We perform Spearman’s rank correlation analysis between receptor densities, as measured by autoradiography across all cortical layers of all ROIs and the abovementioned prediction of mRNA expression, pre-processed microarray data and RNA-Seq data obtained from Allen Human Brain atlas. Correlation coefficient rs was considered low if its absolute value was below 0.5 (Cohen, 1992).

The significance of the correlation test is assessed by the spatial autocorrelation preserving, generative null modeling framework implemented in Brainsmash (Burt et al., 2020). In particular, 10,000 surrogate maps preserving 5 nearest neighbors are created for each transcript and subsequently correlated with respective protein density data. The proportion of the absolute value of correlation coefficients produced by the surrogate maps higher than the absolute value of empirical data calculated correlation coefficient defines p-value (p_SA_). All p-values are corrected for multiple testing using Bonferroni correction for the number of targets and the result is considered as significant when p_sA_ < 0.05. Statistical analyses are performed using MATLAB (R2018b, MathWorks, Natick, USA).

## 3. Results

### 3.1 Predicted mRNA expression

The primary analysis correlating interpolated mRNA expression based on the AHBA to autoradiography data provides a diverse picture. Comparison between mRNA expression and receptor density of kainate receptor and *GRIK1, GRIK2*; GABA_A_ receptor and *GABRA1, GABRA4, GABRA6, GABRB2, GABRG2, GABRG3*; muscarinic acetylcholine receptor M3 and *CRHM3*; adrenoceptors α_1_ and *ADRA1B*; α_2_ and *ADRA2A* and serotonin 1A receptor and *HTR1A* (see Figure 2a) reveal strong positive associations with almost all Spearman’s correlation coefficient above 0.5 (see Table 2). On the other hand, the correlation between levels of mRNA and associated protein abundance as measured by autoradiography unveil significant negative correlations for GABA_A_ receptor and *GABRA3, GABRA5* and *GABRB1* (Table 2), (Figure 2b). Despite a small number of strong links between mRNA transcripts and target receptor distributions, most of the comparisons yield weak non-significant. The overviews of all correlation results are listed in supplementary tables 2-6.

**Table 2:** Gene expressions represented by predicted mRNA patterns significantly (p_SA_ < 0.05) correlated with receptor distributions. Negative correlations are listed at the bottom of the table.

| Receptor | Gene ID | Gene | $r_s$ | $p_{SA}$ |
| --- | --- | --- | --- | --- |
| 5-HT<sub>1A</sub> | *HTR1A* | serotonin receptor 1A | 0.708 | 0.000 |
| GABA<sub>A</sub> | *GABRG2* | GABA<sub>A</sub> receptor subunit $\gamma_2$ | 0.738 | 0.000 |
| GABA<sub>A</sub> | *GABRG3* | GABA<sub>A</sub> receptor subunit $\gamma_3$ | 0.658 | 0.000 |
| GABA<sub>A</sub> | *GABRA4* | GABA<sub>A</sub> receptor subunit $\alpha_4$ | 0.607 | 0.008 |
| GABA<sub>A</sub> | *GABRA6* | GABA<sub>A</sub> receptor subunit $\alpha_6$ | 0.597 | 0.005 |
| GABA<sub>A</sub> | *GABRA1* | GABA<sub>A</sub> receptor subunit $\alpha_1$ | 0.575 | 0.011 |
| GABA<sub>A</sub> | *GABRB2* | GABA<sub>A</sub> receptor subunit $\beta_2$ | 0.532 | 0.029 |
| Kainate | *GRIK1* | Glu ionotropic receptor kainate type subunit 1 | 0.655 | 0.009 |
| Kainate | *GRIK2* | Glu ionotropic receptor kainate type subunit 2 | 0.629 | 0.014 |
| M<sub>3</sub> | *CHRM3* | cholinergic receptor muscarinic 3 | 0.498 | 0.038 |
| $\alpha_1$ | *ADRA1B* | adrenoceptor $\alpha_1B$ | 0.522 | 0.014 |
| $\alpha_2$ | *ADRA2A* | adrenoceptor $\alpha_2A$ | 0.488 | 0.021 |
| GABA<sub>A</sub> | *GABRA3* | GABA<sub>A</sub> receptor subunit $\alpha_3$ | -0.638 | 0.002 |
| GABA<sub>A</sub> | *GABRB1* | GABA<sub>A</sub> receptor subunit $\beta_1$ | -0.577 | 0.006 |
| GABA<sub>A</sub> | *GABRA5* | GABA<sub>A</sub> receptor subunit $\alpha_5$ | -0.565 | 0.020 |

**Figure 2:**
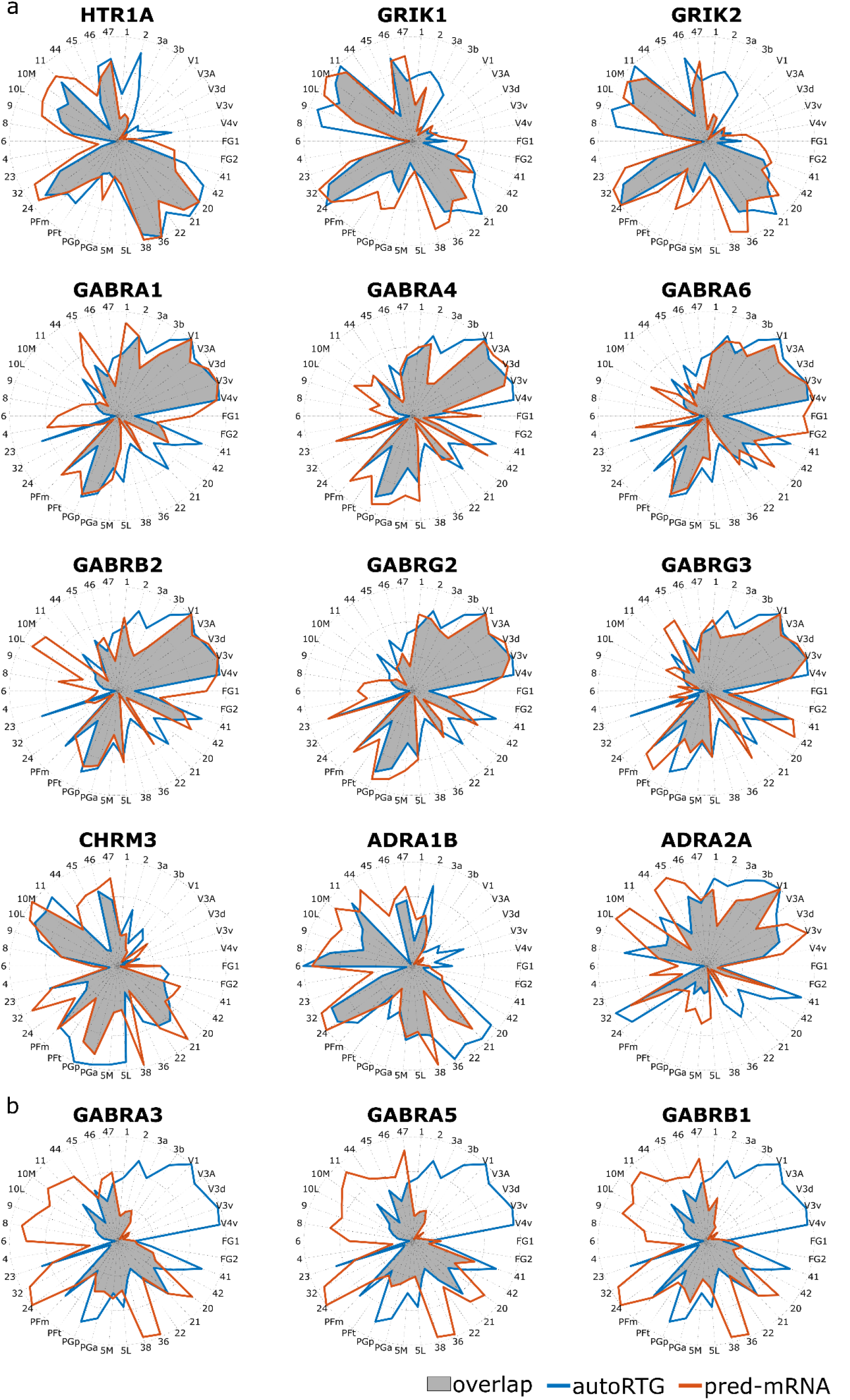
Radar plots showing the relation between interpolated transcriptomic data and autoradiography for significantly a) positive b) negative correlated receptors and gene expressions.

### 3.2 Microarray-based mRNA expression

Complementary analyses, utilizing readily available microarray-based transcriptomic data of the AHBA database, are performed to further investigate the relationship between mRNA expression and protein concentration. Similar to the results with predicted mRNA patterns, we find positive links (Figure 3a) for kainate receptor and *GRIK1, GRIK2*; GABA_A_ receptor and *GABRA1, GABRA4, GABRB2, GABRG2* and serotonin 1A receptor and *HTR1A*, while negative associations (Figure 3b) are unveiled for GABA_A_ receptor and *GABRA3, GABRA5* and *GABRB1*, showing significant strong correlations (Table 3). The majority of the gene expressions show again low and not significant relations with receptor distributions (Supl. table 2-6).

**Figure 3:**
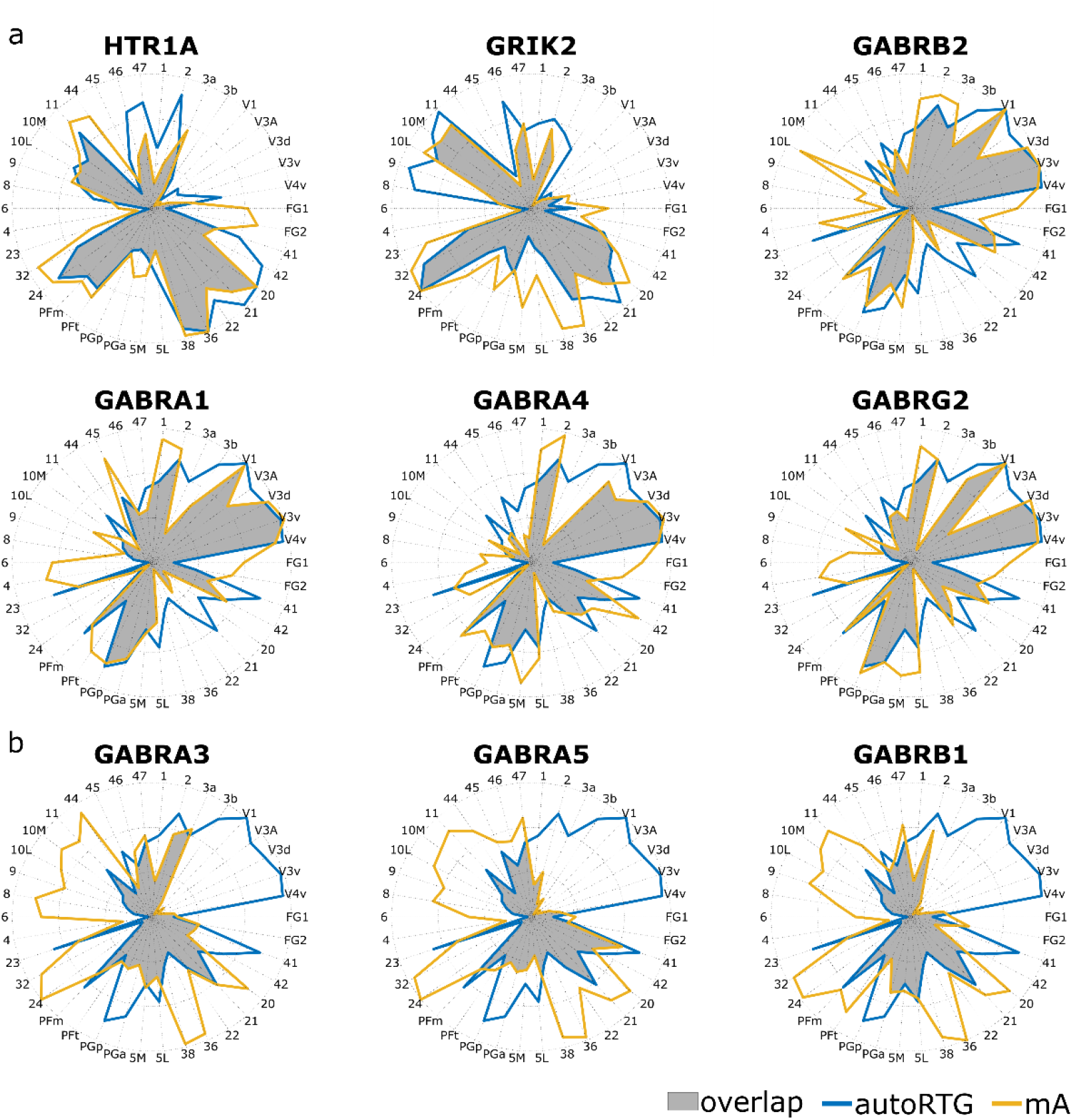
Radar plots showing the relation between microarray transcriptomic data and autoradiography for significantly a) positive b) negative correlated receptors and gene expressions.

**Table 3:**
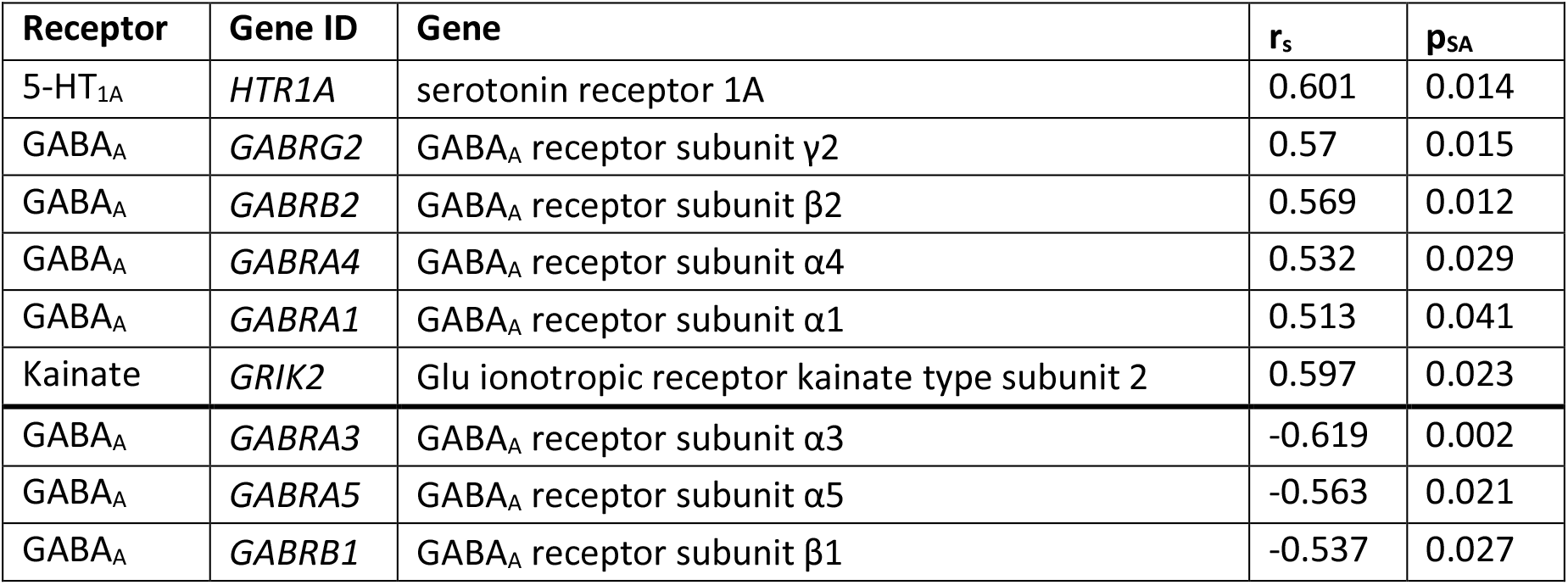
Gene expressions represented by microarray data significantly (p_SA_ < 0.05) correlated with receptor distributions. Negative correlations are listed at the bottom of the table.

| Receptor | Gene ID | Gene | $r_s$ | $p_{SA}$ |
| --- | --- | --- | --- | --- |
| 5-HT<sub>1A</sub> | *HTR1A* | serotonin receptor 1A | 0.601 | 0.014 |
| GABA<sub>A</sub> | *GABRG2* | GABA<sub>A</sub> receptor subunit γ2 | 0.57 | 0.015 |
| GABA<sub>A</sub> | *GABRB2* | GABA<sub>A</sub> receptor subunit β2 | 0.569 | 0.012 |
| GABA<sub>A</sub> | *GABRA4* | GABA<sub>A</sub> receptor subunit α4 | 0.532 | 0.029 |
| GABA<sub>A</sub> | *GABRA1* | GABA<sub>A</sub> receptor subunit α1 | 0.513 | 0.041 |
| Kainate | *GRIK2* | Glu ionotropic receptor kainate type subunit 2 | 0.597 | 0.023 |
| GABA<sub>A</sub> | *GABRA3* | GABA<sub>A</sub> receptor subunit α3 | -0.619 | 0.002 |
| GABA<sub>A</sub> | *GABRA5* | GABA<sub>A</sub> receptor subunit α5 | -0.563 | 0.021 |
| GABA<sub>A</sub> | *GABRB1* | GABA<sub>A</sub> receptor subunit β1 | -0.537 | 0.027 |

### 3.3 RNA-Seq

A further follow-up analysis is looking into the correlation of RNA-Seq data AHBA and target receptor densities. Here, a host of the relations is not significant, yielding only two positive and two negative significant and strong correlations (Table 4). Kainate receptor distribution is positively linked with *GRIK1* as well as GABA_B_ receptor is positively correlated with *GABBR1* (Figure 4a). On the other hand, we find negative associations between adrenoceptor α1 and *ADRA1A* together with GABA_A_ receptor and *GABRQ* (Figure 4b). Overview of the comparison between RNA-Seq and other microarray-based datasets is in the supplementary table 7.

**Figure 4:**
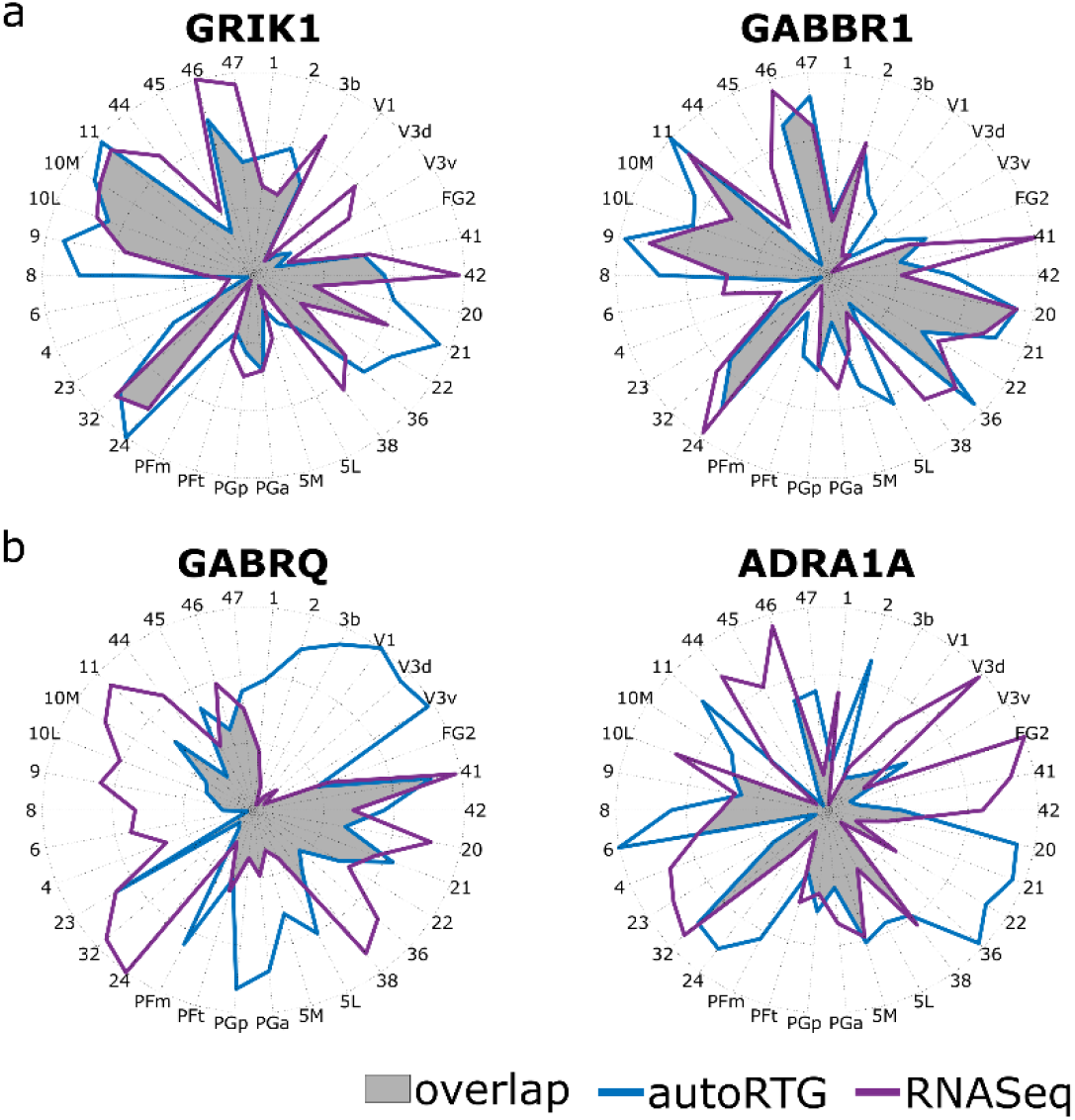
Radar plots showing the relation between RNA-Seq data and autoradiography for significantly a) positive b) negative correlated receptors and gene expressions.

**Table 4:**
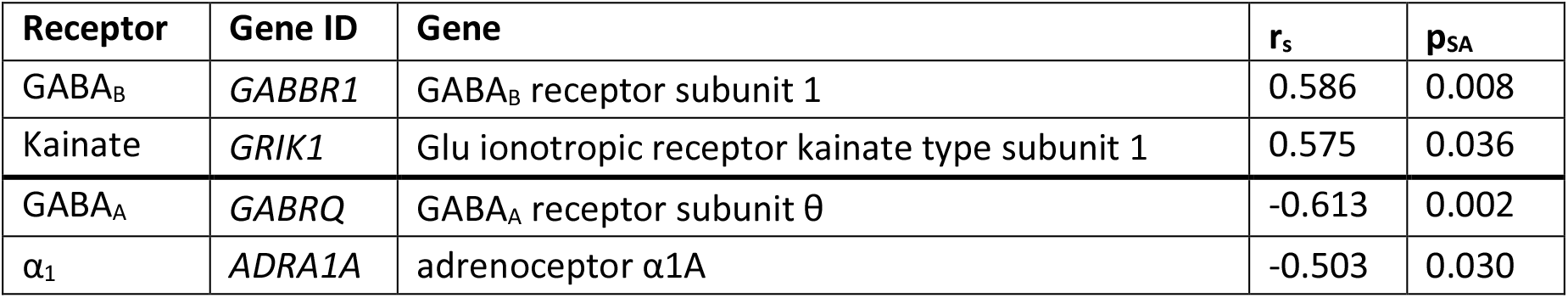
Gene expressions represented RNASeq data significantly (p_SA_ < 0.05) correlated with receptor distributions. Negative correlations are listed at the bottom of the table.

| Receptor | Gene ID | Gene | $r_s$ | $p_{SA}$ |
| --- | --- | --- | --- | --- |
| GABA<sub>B</sub> | *GABBR1* | GABA<sub>B</sub> receptor subunit 1 | 0.586 | 0.008 |
| Kainate | *GRIK1* | Glu ionotropic receptor kainate type subunit 1 | 0.575 | 0.036 |
| GABA<sub>A</sub> | *GABRQ* | GABA<sub>A</sub> receptor subunit $\theta$ | -0.613 | 0.002 |
| $\alpha_1$ | *ADRA1A* | adrenoceptor $\alpha 1A$ | -0.503 | 0.030 |

## 4. Discussion

In this work, we demonstrate associations between mRNA expression and associated protein density patterns for multiple neuroreceptors of the human cerebral cortex. Analysis unveil statistically significant positive correlations across brain areas for some receptors that have been already known to show topological associations with their gene expressions based on PET data, e.g.the serotonin 1 A receptor (Komorowski et al., 2017; Unterholzner et al., 2020). Similarly, kainate receptor showed significant similarities in the spatial distribution of receptor densities and mRNA expression patterns of subunits GluK1 and GluK2 (*GRIK1, GRIK2*). These are allowing for the formation of homomeric receptors and show different pharmacological properties as the rest of the subunits (Fisher and Fisher, 2014). On the other hand, a previously shown strong correlation between gene expression and PET measured receptor distribution of e.g. serotonin 2A receptor (Komorowski et al., 2017) was not confirmed here. This discrepancy might come from the different origin of the proteomic data that is subunit specific for PET, while autoradiography is quantifying the distribution of the whole 5-HT2 receptor.

Counterintuitively, cortical distributions of receptors and their expression patterns also showed a few strong negative relations suggesting a solid, indirect relation underlining the possibility of intricate regulatory circuitry at an intermediate level between mRNA expression and translation. Despite several significant results, most of the comparisons yield weak, statistically not significant correlations between transcriptomic and proteomic information. The results revealed by our analysis fit the outcomes of previous publications correlating PET and mRNA data (Godbersen et al., 2021; Maier et al., 2009)

The differences in correlation observed between variety receptor targets have been attributed to a multitude of factors in the recent past. Foremost is the post-translational modifications (PTMs), which can lead to an apparent local mismatch of mRNA expression to the gene product. The biological need for such modifications arises with the relatively long half-life of eukaryotic mRNA and therefore the need to regulate resulting protein levels in short time frames (Raghavan et al., 2002). This, on a biological level, can be achieved by influencing, for example, translational efficiency and the rate of protein degradation as well as a host of other effects elicited by PTMs. The total extent of influences at different steps in translation may vary significantly between organisms, cell populations and metabolic states where overabundance or lack of available ribosomes might play a decisive role in determining resulting concentrations of the gene product (Hershey et al., 2012). Factors that proved to play a major role here are ribosome transit times (Fan and Penman, 1970) and polysome profiles, which both offer additional information regarding the phase of protein synthesis affected by specific changes. In addition, posttranslational translocation of the proteins may also play an integral role in PTMs affecting the spatial distribution of receptors (Johnson et al., 2013). These scientific undertakings helped reveal that translational control can be influenced on a global level, in a specific subset of mRNA, or on a level of single mRNA. Such global changes can be brought upon by modifications to the overall translational processes while even more specific changes can be mediated by microRNA or trans-acting proteins (Gebauer et al., 2012).

Furthermore, individual subunit transcripts might not always offer the same quality of surrogate for receptor distribution. A prime example for this is the GABA_A_ receptor, which is a complex pentamer usually made up of distinct subunit, α(1–6), β(1–4), γ(1–3), δ, ε, θ, π and ρ(1–3). The most abundant isomer in the human cerebral cortex is typically comprised of two α1, two β2 and one γ2 subunit(s) (Macdonald and Olsen, 1994; Olsen and Sieghart, 2008). Next to this most frequent pentamer, multiple other isoforms have been detected (Farrar et al., 1999; Tretter et al., 1997). Noteworthy, gene expression patterns of all three aforementioned subunits show a strong, positive significant correlation with GABA_A_ receptor distribution. While distinct variants make up most of the regional receptor populations, parts of the brain show different expression patterns leaving α1β2γ2 isomer as an exception rather than the rule. Namely, several GABA_A_ subunits (α3, α5, β1) show significantly negative correlations between receptor distribution and its gene expressions. This discrepancy could be possibly attributed to the most abundant subunits that might mask the real correlation between receptor subunits and their gene expression. Besides subtype variants, for GABA-receptors extensive post-translational modifications have been demonstrated offering an additional venue for effective decoupling of mRNA expression and resulting protein concentration (Campbell and Tyagarajan, 2019). The origin of unexpected low correlation values might also lie on the side of the autoradiography dataset. Previous studies have already proven that concentrations of endogenous neurotransmitters can significantly affect the radioligand binding to its target suggesting that measured protein distribution might be not representative (Sasaki et al., 2002). Moreover, as previously suggested for the GABA_A_ or 5-HT_2_ receptors, the masking effect of the most abundant subunits and subtypes might affect the correlation. For example, [^3^H]-Epibatidine used for labeling nicotinic α_4_β_2_ receptor is an agonist for subunits α_2_-α_7_ binding to all of them with similar affinity (Gaulton et al., 2017). In particular, alongside the α_4_β_2_ receptor, the homomeric α_7_ receptor which is also widely present across the brain (Gotti et al., 2006), might cause masking of the real α4 distribution. On the other hand [^3^H]8-OH-DPAT, a highly specific ligand for 5-HT_1A_ receptor has a lower affinity to other 5-HT_1_ subtypes (Newman-Tancredi et al., 1998), which is further supported by the strong correlation between its spatial distribution and gene expression. These discrepancies in the binding of ligands suggest, that poor correlation between most of the receptor distribution and gene expression patterns might be affected by different affinities to individual receptor subunits.

To fully grasp the picture of protein abundance resulting from mRNA expression patterns, it is imperative to evaluate not only total protein and mRNA in corresponding samples, ideally from the same donor, but to also quantify each step of protein synthesis along the way. This will require significant effort, as methods for global identification and quantification of both transcriptome and resulting translatome still require a high level of expertise in experimental design and methodical calculations (Arnatkevičiūtė et al., 2019). Nowadays, most of microarray data limitations might be overcome by employing next generation sequencing data (Stark et al., 2019; Wang et al., 2009), enabling high throughput parallel sequence detection enabling procedural counting of transcripts to allow for direct quantification. However, using additional data set of RNA-Seq data for comparison to receptor distributions confirmed only a couple of results coming from microarray data analysis (e.g. for kainate receptor subunit 1 or GABA_A_ receptor subunit α5) while most of the results were not significant which can be accounted to rather scarce RNA-Seq data set of AHBA. The limited information contained in the RNA-Seq AHBA data is further upheld by the counterintuitive results of the comparison to microarray data sets, showing a rather weak correlation for most of the investigated gene expressions highlight for more comprehensive RNA-Seq database. Ultimately, methods like span single cell *in vivo* experiments and high throughput sequencing experiments for mRNA might provide promising quantitative information to further uncover underlying links between receptor distributions and mRNA levels.

### 4.1 Limitations

Firstly, besides the commonly known limitation of Allen Human Brain atlas genetic data sets (most of the sparsely distributed samples come from 6 left hemispheres while only 2 right hemispheres are available), the autoradiography data set stems from a different set of donors than the transcriptomic data. While we can partially account for the Allen Human Brain atlas limitations by mirroring the samples along the sagittal plane and using the predicted cortical mRNA expression patterns (Gryglewski et al., 2020), the inter-subject variability between transcriptomic and proteomic data sources could not be resolved. Additionally, at this point, it appears that the problem of subdividing used autoradiographic signals to the corresponding receptor subtypes remains unsolvable, without further measurements using subunit-specific tracers. It is known that subtypes of neuro-receptors exhibit different binding potentials – therefore, it cannot be ruled out in these cases, that the majority of the signal is due to a single variant, subunit or conformation, which confers way stronger binding than its counterparts. This fact diminishes the overall explanatory power of the analysis at hand but also illustrates the need for more advanced models to allow for the estimation of specific protein abundancy.

## 5. Conclusion

In the present work, we report a comparison of available high-quality data characterizing cortical receptor distribution at the level of mRNA expression as well as protein abundance. We identified several positive as well as negative links between the scrutinized modalities, however, most of the comparisons showed no direct relation between protein density and transcriptomic information.

Current results show that gene expression should not be directly used as a surrogate for protein distribution rather as additional information in multimodal studies to further investigate the effect of numerous processes happening on the way from mRNA to protein. The role of posttranscriptional processes and posttranslational modification cannot be ignored and should motivate further studies on potential mRNA markers associated with them.

## 6. Data availability statement

Due to data protection laws, processed data is available from the authors upon reasonable request. Please contact with any questions or requests.

## 7. Acknowledgements

M. Murgaš is funded by the Austrian Science Fund FWF, DOC 33-B27. MB. Reed is a recipient of a DOC fellowship of the Austrian Academy of Sciences at the Department of Psychiatry and Psychotherapy, Medical University of Vienna. We would like to thank Prof. Zilles (†), Prof. Amunts and Dr. Palomero-Gallagher of the Research Centre Jülich, Institute of Neuroscience and Medicine (INM-1), Jülich, Germany, (Zilles and Palomero-Gallagher, 2017), as well as the Allen Institute for Brain Science, Seattle, USA (Hawrylycz et al., 2012) for making their valuable data sets publicly available.

