## Supplementary material for "Correlation of receptor density and mRNA expression patterns in the human cerebral cortex": Supplemetary Material

### Supplementary Information

| Area ID in Julich-Brain | Brodmann Area ID | Description |
|---|---|---|
| 1 | 1 | primary somatosensory cortex |
| 2 | 2 | primary somatosensory cortex |
| 3a | 3 | primary somatosensory cortex |
| 3b | 3 | primary somatosensory cortex |
| 4 | 4 | primary motor cortex |
| 5L | 5 | area of the higher unimodal somatosensory cortex in superior parietal cortex |
| 5M | 5 | area of the higher unimodal somatosensory cortex in superior parietal cortex |
| 6 | 6 | premotor cortex |
| 8 | 8 | isocortical prefrontal area |
| 9 | 9 | isocortical prefrontal area |
| 10L | 10 | isocortical prefrontal area |
| 10M | 10 | isocortical prefrontal area |
| 11 | 11 | isocortical prefrontal area |
| V1 | 17 | primary visual cortex |
| V3A | 19 | higher visual cortical area |
| V3d | 19 | higher visual cortical area |
| V3v | 19 | higher visual cortical area |
| V4v | 19 | higher visual cortical area |
| FG1 | - | fusiform gyrus of higher visual area |
| FG2 | - | fusiform gyrus of higher visual area |
| 20 | 20 | multimodal temporal area |
| 21 | 21 | multimodal temporal area |
| 22 | 22 | primary unimodal auditory area |
| 23 | 23 | isocortical cingulate area |
| 24 | 24 | periarchicortical cingulate area |
| 32 | 32 | isocortical cingulate area |
| 36 | 36 | multimodal temporal area |
| 38 | 38 | multimodal temporal area |
| PGa | 39 | inferior parietal area, supramarginal gyrus of BA39 |
| PGp | 39 | inferior parietal area, supramarginal gyrus of BA39 |
| PFt | 40 | inferior parietal area, angular gyrus of BA40 |
| PFm | 40 | inferior parietal area, angular gyrus of BA40 |
| 41 | 41 | higher unimodal auditory area |
| 42 | 42 | higher unimodal auditory area |
| 44 | 44 | ventral part of BA44 in Broca's region |
| 45 | 45 | anterior part of BA45 in Broca's region |
| 46 | 46 | isocortical prefrontal area |
| 47 | 47 | isocortical prefrontal area |

*Suppl. table 1: Overveiw of cytoarchitectonic regions used for the analysis taken from Julich-Brain atlas presented in* (Amunts et al., 2020).

| Target (Receptor) | Gene Symbol | pred-mRNA | | mA | | RNASeq | |
|---|---|---|---|---|---|---|---|
| | | $r_s$ | $p_{SA}$ | $r_s$ | $p_{SA}$ | $r_s$ | $p_{SA}$ |
| **AMPA** | ***GRIA1*** | 0.304 | 1.000 | 0.365 | 0.995 | 0.522 | 0.088 |
| **AMPA** | ***GRIA2*** | 0.227 | 1.000 | -0.169 | 1.000 | 0.384 | 0.820 |
| **AMPA** | ***GRIA3*** | -0.450 | 0.290 | -0.147 | 1.000 | 0.364 | 0.976 |
| **AMPA** | ***GRIA4*** | 0.272 | 1.000 | 0.218 | 1.000 | 0.199 | 1.000 |
| **NMDA** | ***GRIN1*** | -0.338 | 1.000 | 0.573 | 1.000 | -0.268 | 1.000 |
| **NMDA** | ***GRIN2A*** | 0.055 | 1.000 | 0.103 | 1.000 | -0.137 | 1.000 |
| **NMDA** | ***GRIN2B*** | -0.018 | 1.000 | 0.102 | 1.000 | -0.014 | 1.000 |
| **NMDA** | ***GRIN2C*** | -0.127 | 1.000 | 0.086 | 1.000 | 0.204 | 0.480 |
| **NMDA** | ***GRIN2D*** | -0.317 | 1.000 | 0.110 | 1.000 | 0.404 | 1.000 |
| **NMDA** | ***GRIN3A*** | -0.199 | 1.000 | -0.164 | 1.000 | -0.003 | 1.000 |
| **NMDA** | ***GRIN3B*** | -0.054 | 1.000 | 0.045 | 1.000 | -0.004 | 1.000 |
| **kainate** | ***GRIK1*** | 0.655 | 0.009 | 0.297 | 1.000 | 0.575 | 0.036 |
| **kainate** | ***GRIK2*** | 0.629 | 0.014 | 0.597 | 0.023 | 0.564 | 0.058 |
| **kainate** | ***GRIK3*** | 0.497 | 0.168 | 0.412 | 0.585 | 0.433 | 0.509 |
| **kainate** | ***GRIK4*** | 0.271 | 1.000 | 0.113 | 1.000 | 0.113 | 1.000 |
| **kainate** | ***GRIK5*** | 0.351 | 1.000 | 0.379 | 0.836 | 0.529 | 0.083 |

*Suppl. table 2: Correlation coefficients of predicted mRNA expression (pred-mRNA), microarray (mA) and RNA Sequencing (RNA-Seq) data to the autoradiographic data representing AMPA, NMDA and kainate ionotropic glutamate receptors.*

| Target (Receptor) | Gene Symbol | pred-mRNA | | mA | | RNASeq | |
|---|---|---|---|---|---|---|---|
| | | $r_s$ | $p_{SA}$ | $r_s$ | $p_{SA}$ | $r_s$ | $p_{SA}$ |
| $\mathbf{M_1}$ | ***CHRM1*** | 0.117 | 1.000 | -0.315 | 1.000 | 0.101 | 1.000 |
| $\mathbf{M_2}$ | ***CHRM2*** | 0.279 | 1.000 | -0.253 | 1.000 | 0.249 | 1.000 |
| $\mathbf{M_3}$ | ***CHRM3*** | 0.498 | 0.038 | 0.415 | 0.180 | 0.132 | 1.000 |
| $\boldsymbol{\alpha_4\beta_2}$ | ***CHRNA4*** | -0.228 | 1.000 | -0.025 | 1.000 | 0.200 | 0.971 |
| $\boldsymbol{\alpha_4\beta_2}$ | ***CHRNB2*** | -0.007 | 1.000 | 0.050 | 1.000 | 0.300 | 1.000 |

*Suppl. table 3: Correlation coefficients of predicted mRNA expression (pred-mRNA), microarray (mA) and RNA Sequencing (RNA-Seq) data to the autoradiographic data of acetylcholine receptors muscarinic $M_1$, $M_2$, $M_3$ and nicotinic $\alpha_4\beta_2$.*

| Target (Receptor) | Gene Symbol | pred-mRNA | | mA | | RNASeq | |
|---|---|---|---|---|---|---|---|
| | | $r_s$ | $p_{SA}$ | $r_s$ | $p_{SA}$ | $r_s$ | $p_{SA}$ |
| $\mathbf{GABA_A}$ | ***GABRA1*** | 0.575 | 0.00 | 0.513 | 0.00 | 0.043 | 0.81 |
| $\mathbf{GABA_A}$ | ***GABRA2*** | -0.387 | 0.02 | -0.254 | 0.12 | -0.341 | 0.04 |
| $\mathbf{GABA_A}$ | ***GABRA3*** | -0.638 | 0.00 | -0.619 | 0.00 | -0.487 | 0.00 |
| $\mathbf{GABA_A}$ | ***GABRA4*** | 0.607 | 0.00 | 0.532 | 0.00 | -0.072 | 0.69 |
| $\mathbf{GABA_A}$ | ***GABRA5*** | -0.565 | 0.00 | -0.563 | 0.00 | -0.407 | 0.02 |
| $\mathbf{GABA_A}$ | ***GABRA6*** | 0.597 | 0.00 | -0.041 | 0.81 | 0.195 | 0.26 |
| $\mathbf{GABA_A}$ | ***GABRB1*** | -0.577 | 0.00 | -0.537 | 0.00 | -0.355 | 0.04 |
| $\mathbf{GABA_A}$ | ***GABRB2*** | 0.532 | 0.00 | 0.569 | 0.00 | 0.008 | 0.97 |
| $\mathbf{GABA_A}$ | ***GABRB3*** | -0.467 | 0.00 | -0.502 | 0.00 | -0.351 | 0.04 |
| $\mathbf{GABA_A}$ | ***GABRD*** | 0.461 | 0.00 | 0.415 | 0.01 | 0.102 | 0.56 |
| $\mathbf{GABA_A}$ | ***GABRE*** | -0.175 | 0.29 | -0.161 | 0.34 | 0.268 | 0.13 |
| $\mathbf{GABA_A}$ | ***GABRG1*** | -0.358 | 0.03 | -0.168 | 0.31 | 0.131 | 0.46 |
| $\mathbf{GABA_A}$ | ***GABRG2*** | 0.738 | 0.00 | 0.570 | 0.00 | -0.010 | 0.95 |
| $\mathbf{GABA_A}$ | ***GABRG3*** | 0.658 | 0.00 | 0.183 | 0.27 | -0.172 | 0.32 |

| | | | | | | | |
|---|---|---|---|---|---|---|---|
| **GABA$_{A}$** | ***GABRP*** | -0.072 | 0.67 | -0.097 | 0.57 | -0.277 | 0.11 |
| **GABA$_{A}$** | ***GABRQ*** | -0.504 | 0.00 | -0.250 | 0.22 | -0.613 | 0.00 |
| **GABA$_{A/BZ}$** | ***GABRA1*** | 0.113 | 0.02 | 0.151 | 0.00 | 0.586 | 0.00 |
| **GABA$_{A/BZ}$** | ***GABRA2*** | 0.224 | 0.27 | 0.151 | 0.74 | 0.119 | 0.51 |
| **GABA$_{A/BZ}$** | ***GABRA3*** | -0.085 | 0.50 | -0.168 | 0.37 | -0.140 | 0.42 |
| **GABA$_{A/BZ}$** | ***GABRA5*** | -0.115 | 0.61 | -0.110 | 0.31 | -0.261 | 0.14 |
| **GABA$_{A/BZ}$** | ***GABRB1*** | -0.020 | 0.49 | 0.079 | 0.52 | -0.283 | 0.10 |
| **GABA$_{A/BZ}$** | ***GABRB2*** | 0.286 | 0.64 | 0.170 | 0.35 | 0.213 | 0.22 |
| **GABA$_{A/BZ}$** | ***GABRB3*** | 0.015 | 0.91 | -0.050 | 0.64 | -0.258 | 0.14 |
| **GABA$_{A/BZ}$** | ***GABRD*** | 0.080 | 0.08 | 0.112 | 0.30 | -0.076 | 0.67 |
| **GABA$_{A/BZ}$** | ***GABRE*** | 0.046 | 0.92 | 0.039 | 0.76 | -0.321 | 0.07 |
| **GABA$_{A/BZ}$** | ***GABRG1*** | 0.166 | 0.63 | 0.241 | 0.50 | 0.138 | 0.44 |
| **GABA$_{A/BZ}$** | ***GABRG2*** | 0.385 | 0.78 | 0.282 | 0.82 | 0.119 | 0.50 |
| **GABA$_{A/BZ}$** | ***GABRG3*** | 0.281 | 0.31 | 0.425 | 0.14 | 0.033 | 0.85 |
| **GABA$_{A/BZ}$** | ***GABRP*** | -0.152 | 0.02 | -0.023 | 0.09 | 0.011 | 0.95 |
| **GABA$_{A/BZ}$** | ***GABRQ*** | -0.027 | 0.09 | -0.109 | 0.01 | -0.323 | 0.06 |
| **GABA$_{B}$** | ***GABBR1*** | 0.390 | 0.36 | 0.464 | 0.90 | -0.101 | 0.57 |
| **GABA$_{B}$** | ***GABBR2*** | -0.182 | 0.87 | 0.055 | 0.60 | -0.353 | 0.04 |

*Suppl. table 4: Correlation coefficients of predicted mRNA expression (pred-mRNA), microarray (mA) and RNA Sequencing (RNA-Seq) data to the autoradiographic data of GABA$_{A}$, GABA$_{A}$ benzodiazepine binding site and GABA$_{B}$ receptors.*

| **Target (Receptor)** | **Gene Symbol** | **pred-mRNA** | | **mA** | | **RNASeq** | |
|---|---|---|---|---|---|---|---|
| | | $r_s$ | $p_{SA}$ | $r_s$ | $p_{SA}$ | $r_s$ | $p_{SA}$ |
| **$\alpha_1$** | ***ADRA1A*** | -0.028 | 1.000 | -0.131 | 1.000 | -0.503 | 0.030 |
| **$\alpha_1$** | ***ADRA1B*** | 0.522 | 0.014 | 0.191 | 1.000 | 0.284 | 1.000 |
| **$\alpha_1$** | ***ADRA1D*** | -0.147 | 1.000 | -0.256 | 1.000 | 0.060 | 1.000 |
| **$\alpha_2$** | ***ADRA2A*** | 0.488 | 0.021 | 0.281 | 1.000 | 0.204 | 1.000 |
| **$\alpha_2$** | ***ADRA2B*** | 0.137 | 1.000 | -0.256 | 1.000 | -0.207 | 1.000 |
| **$\alpha_2$** | ***ADRA2C*** | -0.195 | 1.000 | -0.264 | 1.000 | 0.012 | 1.000 |

*Suppl. table 5: Correlation coefficients of predicted mRNA expression (pred-mRNA), microarray (mA) and RNA Sequencing (RNA-Seq) data to the autoradiographic data of noradrenaline receptors $\alpha_1$ and $\alpha_2$.*

| **Target (Receptor)** | **Gene Symbol** | **pred-mRNA** | | **mA** | | **RNASeq** | |
|---|---|---|---|---|---|---|---|
| | | $r_s$ | $p_{SA}$ | $r_s$ | $p_{SA}$ | $r_s$ | $p_{SA}$ |
| **5-HT$_{1A}$** | ***HTR1A*** | 0.708 | 0.000 | 0.601 | 0.014 | 0.544 | 0.056 |
| **5-HT$_{2}$** | ***HTR2A*** | 0.178 | 1.000 | 0.164 | 1.000 | -0.362 | 0.430 |
| **5-HT$_{2}$** | ***HTR2B*** | -0.265 | 1.000 | 0.065 | 1.000 | -0.062 | 1.000 |
| **5-HT$_{2}$** | ***HTR2C*** | 0.150 | 1.000 | 0.205 | 1.000 | 0.057 | 1.000 |
| **D$_{1}$** | ***DRD1*** | 0.348 | 0.641 | 0.345 | 0.639 | -0.128 | 1.000 |

*Suppl. table 6: Correlation coefficients of predicted mRNA expression (pred-mRNA), microarray (mA) and RNA Sequencing (RNA-Seq) data to the autoradiographic data of serotonin and dopamine receptors.*

| Gene ID | Gene Name | $r_s$ (vs mRNA) | $r_s$ (vs mA) |
|---|---|---|---|
| *GRIA1* | Glu ionotropic receptor AMPA type subunit 1 | 0.376 | 0.695 |
| *GRIA2* | Glu ionotropic receptor AMPA type subunit 2 | 0.045 | -0.287 |
| *GRIA3* | Glu ionotropic receptor AMPA type subunit 3 | -0.151 | -0.306 |
| *GRIA4* | Glu ionotropic receptor AMPA type subunit 4 | 0.169 | 0.143 |
| *GRIN1* | Glu ionotropic receptor NMDA type subunit 1 | 0.309 | 0.021 |
| *GRIN2A* | Glu ionotropic receptor NMDA type subunit 2A | 0.327 | 0.217 |
| *GRIN2B* | Glu ionotropic receptor NMDA type subunit 2B | 0.473 | 0.224 |
| *GRIN2C* | Glu ionotropic receptor NMDA type subunit 2C | -0.003 | -0.133 |
| *GRIN2D* | Glu ionotropic receptor NMDA type subunit 2D | -0.026 | 0.081 |
| *GRIN3A* | Glu ionotropic receptor NMDA type subunit 3A | 0.845 | 0.663 |
| *GRIN3B* | Glu ionotropic receptor NMDA type subunit 3B | 0.109 | -0.010 |
| *GRIK1* | Glu ionotropic receptor kainate type subunit 1 | 0.586 | 0.274 |
| *GRIK2* | Glu ionotropic receptor kainate type subunit 2 | 0.720 | 0.609 |
| *GRIK3* | Glu ionotropic receptor kainate type subunit 3 | 0.531 | 0.422 |
| *GRIK4* | Glu ionotropic receptor kainate type subunit 4 | 0.525 | 0.342 |
| *GRIK5* | Glu ionotropic receptor kainate type subunit 5 | 0.469 | 0.389 |
| *GABRA1* | $GABA_A$ receptor subunit α1 | 0.163 | 0.077 |
| *GABRA2* | $GABA_A$ receptor subunit α2 | 0.635 | 0.278 |
| *GABRA3* | $GABA_A$ receptor subunit α3 | 0.639 | 0.602 |
| *GABRA4* | $GABA_A$ receptor subunit α4 | -0.145 | -0.370 |
| *GABRA5* | $GABA_A$ receptor subunit α5 | 0.886 | 0.888 |
| *GABRA6* | $GABA_A$ receptor subunit α6 | 0.399 | 0.234 |
| *GABRB1* | $GABA_A$ receptor subunit β1 | 0.577 | 0.441 |
| *GABRB2* | $GABA_A$ receptor subunit β2 | 0.283 | 0.110 |
| *GABRB3* | $GABA_A$ receptor subunit β3 | 0.831 | 0.685 |
| *GABRD* | $GABA_A$ receptor subunit δ | 0.583 | 0.598 |
| *GABRE* | $GABA_A$ receptor subunit Ɛ | 0.614 | 0.728 |
| *GABRG1* | $GABA_A$ receptor subunit γ1 | 0.448 | 0.432 |
| *GABRG2* | $GABA_A$ receptor subunit γ2 | 0.067 | 0.089 |
| *GABRG3* | $GABA_A$ receptor subunit γ3 | -0.048 | -0.266 |
| *GABRP* | $GABA_A$ receptor subunit π | -0.159 | -0.091 |
| *GABRQ* | $GABA_A$ receptor subunit θ | 0.829 | 0.176 |
| *GABBR1* | $GABA_B$ receptor subunit 1 | 0.556 | 0.470 |
| *GABBR2* | $GABA_B$ receptor subunit 2 | 0.160 | 0.119 |
| *CHRM1* | cholinergic receptor muscarinic 1 | 0.336 | 0.240 |
| *CHRM2* | cholinergic receptor muscarinic 2 | 0.055 | -0.183 |
| *CHRM3* | cholinergic receptor muscarinic 3 | 0.444 | 0.346 |
| *CHRNA4* | cholinergic receptor nicotinic α4 subunit | 0.108 | 0.179 |
| *CHRNB2* | cholinergic receptor nicotinic β2 subunit | -0.098 | -0.121 |
| *ADRA1A* | adrenoceptor α1A | -0.165 | 0.065 |
| *ADRA1B* | adrenoceptor α1B | 0.374 | -0.092 |
| *ADRA1D* | adrenoceptor α1C | 0.383 | 0.234 |
| *ADRA2A* | adrenoceptor α2A | 0.400 | 0.386 |
| *ADRA2B* | adrenoceptor α2B | -0.206 | -0.293 |
| *ADRA2C* | adrenoceptor α2C | 0.149 | 0.400 |
| *HTR1A* | serotonin receptor 1A | 0.727 | 0.670 |
| *HTR2A* | serotonin receptor 2A | 0.104 | 0.052 |
| *HTR2B* | serotonin receptor 2B | 0.004 | 0.228 |
| *HTR2C* | serotonin receptor 2C | 0.830 | 0.658 |
| *DRD1* | dopamine receptor D1 | -0.001 | -0.374 |

*Suppl. table 7: Overview of correlation coefficients between RNA-Seq data and predicted mRNA expression (3rd column), microarray data (4th column).*
